# New insights into tomato CLE peptide repertoire and perception mechanisms

**DOI:** 10.1101/2022.01.21.477294

**Authors:** Samy Carbonnel, Laurent Falquet, Ora Hazak

## Abstract

Precision in sensing the environmental cues and adjusting the growth and the physiology of the root system are necessary for plant robustness. Plants achieve their phenotypic plasticity by tightly controlling and buffering developmental decisions. In addition to the classical plant hormones that mediate plant development and stress responses, the CLE peptides constitute an additional crucial level of regulation. While the CLV3-CLV1 module appears to be highly conserved to control the proliferation of the shoot apical meristem stem cells, we do not yet fully understand the function of the additional *CLE* genes and whether they act in a similar way across the plant species, including tomato. Due to the small gene size and high sequence variability, it is extremely difficult to precisely annotate *CLE* genes in plant genomes. Here we present our analysis of the *CLE* family in tomato, based on a combination of iterative tBLASTn and Hidden-Markov-Model (HMM), which allowed us to identify thirty-seven new *SlCLE*s in addition to the fifteen reported previously. We could confirm the biological activities of selected SlCLEs in suppressing root meristematic cell divisions. We show that root response is mediated by *SlCLAVATA2*, indicating the conservation of CLE perception mechanism.

**One-sentence summary:** Using a combination of iterative tBLASTn and Hidden-Markov-Model approaches, we uncovered 37 new tomato *CLE* genes predominantly expressed in roots, and we showed a conserved effect on root meristem arrest, that was *SlCLAVATA2*-dependent.

**Highlights:**

- We applied a combined approach of iterative tBLASTn and Hidden-Markov-Model to identify fifty-two tomato *SlCLE* genes, including thirty-seven new genes
- All identified genes encode for pre-propeptides with a single CLE-domain containing conserved residues similar to Arabidopsis
- Analyzing the publicly available RNAseq datasets, we could confirm the expression of *SlCLE* genes that was often associated specifically with root or shoot, a certain developmental stage of the fruit, or with drought stress conditions
- Remarkably, the majority of *SlCLE* genes are predominantly expressed in the root tissues
- We showed the conserved inhibitory effect on the root meristem and columella cells division for the selected SlCLE peptides that were *SlCLAVATA2*-dependent.

## 1. Introduction

Plant roots explore the soil for water and minerals that are distributed irregularly in space and time. They grow towards the resources and limit the growth towards the unfavorable substances [1]. Remarkably, roots integrate the inputs from the soil environment, to adjust the growth and to maximize the nutrient and water uptake. The biology of the root growth and responses to the environment is an emerging topic [2] and therefore, finding new regulators and pathways that underlie the root growth under stressful conditions is extremely important. In the last decades, the Arabidopsis research yielded many key players that shape the root system, including plant hormones, small RNAs, transcription factors, and small signaling peptides [3, 4]. At the same time, we still do not know whether the mechanisms observed in Arabidopsis are fully conserved in the eudicot and the monocot crop species.

One of the most studied groups of hormone-like peptides that have been shown to mediate root vascular development, root stem cell niche maintenance, and responses to the availability of water, phosphate, nitrate in the soil, sugar levels, and root-to-shoot communication is the CLAVATA3/EMBRYO-SURROUNDING REGION-RELATED (CLE) family [5-12]. The CLE genes are relatively small and encode for non-functional pre-propeptides of about 100 amino acids. To become active peptides, additional processing, including cleavage by subtilases [13], and, often prolines hydroxylation and glycosylation are necessary[13-16]. CLE peptides are secreted to the apoplast, where they are perceived by the Class XI of the leucine-rich repeats receptor-like kinases (LRR-RLKs) [9, 17]. In Arabidopsis, in addition to the CLAVATA1 receptor, three BARELY ANY MERISTEM (BAM) receptors have been shown to bind CLE peptides. These receptors have three domains: an extracellular domain, which is responsible for the binding of the ligand, a transmembrane domain, which anchors the receptor in the plasma membrane, and a cytoplasmic kinase domain, that triggers the intracellular signaling by phosphorylating downstream targets. Moreover, the receptor-like kinases CLV3 INSENSITIVE KINASES (CIKs) act as co-receptors both in perceiving the root-active CLE peptides and in the CLV3 signaling in the shoot apical meristem [18, 19]. In addition to these cognate receptors, it has been shown, that also LRR receptor-like protein (LRR-RLP) named CLAVATA2 (CLV2) creates a dimer with the pseudo-kinase CORYNE (CRN) to perceive the full range of CLE peptides in the Arabidopsis root [9].

The unique roles of CLE peptides in mediating shoot and root growth and adaptation to environmental stresses underlies the special interest of the scientific community to study this group of ancient and highly conserved plant peptide hormones not only in model plants but also in crop species. The genome-wide analyses of *CLE* genes have been performed in many plant genomes, including tomato, rice, wheat, maize, soybean, grape, potato and cucumber [20-23]. Due to the small gene size and high sequence variability, the annotation is challenging. It has been previously reported, that in the tomato genome there are only 15 CLE genes [23]. Among them, *SlCLV3* and *SlCLE9* encode for signaling peptides that need to be arabinosylated to control the stem cell proliferation and shoot apical meristem size. Remarkably, the tomato domestication mutation *fascinated (fas)* that led to the increased fruit size, is a result of disruption of the *SlCLV3* promoter that led to the reduction in the gene expression [24]. The *SlCLE9* is the closest paralog of *SlCLV3* and can actively compensate for the absence of *SlCLV3* to buffer the impact on the stem cell niche [24, 25]. The unraveling of additional tomato *CLE* genes and more careful phylogenetic analysis is necessary to fully understand the role of these conserved ligands in tomato development and adaptation to the changing environment.

## 2. Materials and methods

### 2.1 *SlCLE*s identification

#### 2.1.1 Iterative tBLASTn

All previously described *Arabidopsis thaliana* CLE full-length protein (pre-propeptide) sequences were used as queries to search by tBLASTn in *Solanum lycopersicum* genome SL3.0 and SL4.0 in the plant section of the EnsemblGenome database [26]. The hits were then used to search by BLASTp in closely related species of the Solanaceae family (*Nicotiana attenuata, Solanum tuberosum*, and *Capsicum annuum*). The newly identified CLE proteins were exploited to identify by tBLASTn additional similar sequences in tomato’s genome, which were then used to search again in the above Solanaceae-species genomes. Between each iteration, candidate loci were individually confirmed based on the CLE domain sequence and the presence of a signal peptide sequence in 5’.

#### 2.1.2 Hidden-Markov-Model approach

A list of 256 CLE proteins obtained in multiple species (*A. thaliana, N. attenuata, S. tuberosum, Populus trichocarpa, Medicago truncatula, Brachypodium distachyon*, and tomato sequences found in 2.1.1) was aligned with MEGA X [27] and used to build an HMM with HMMER3 [28]. The HMM was used to search *S. lycopersicum* SL4.0 genome with Genewise [29] (the genome was split in chunks of 9 million bp with EMBOSS splitter & seqretsplit [30]). This led to a list of 61 CLE candidates that was concatenated with the 40 CLE of 2.1.1 above. After manual cleaning and removing duplicates, we confirmed a clean list of 57 *CLE* candidates.

#### 2.1.3 Candidate verification

The gene structure of the 57 candidate *CLE* was verified by tBLASTn and BLAT [31] against the SL3.0 genome as in 2.1.1 and by manual evaluation of the resulting hits for the correctness of their exon-intron structure. Five pseudogenes could be identified (with in-frame stop codons or no initiator methionine), leaving a final list of 52 *CLE* genes.

### 2.2 Transcriptomic analysis

We selected four publicly available RNAseq and TRAPseq datasets to search for expression clues of the CLE genes in various tissue types of *S*.*lycopersicum* M82: RNAseqA [32], RNAseqD [33], RNAseqF1 and RNAseqF2 [34], TRAPseq [35].

The selected samples of all the four datasets were remapped to the SL4.0 genome assembly with bwa [36] and samtools [37] to obtain sorted bam files. A Bed file containing the CLE gene positions was created (**CLEgene.bed**) and used to count the reads per gene with bedtools multicov [38]. A heatmap of the logTPM (transcripts per million) for CLE genes counts over all genes was created with a custom-made R script (**script**) for each dataset.

### 2.3 Phylogenetic analysis

Alignments of the CLE proteins sequences were performed in MEGA X [27], using ClustalW (Figure 2) or MUSCLE (Figure S1-S2), and manually corrected. The phylogenetic trees were generated by IQTREE with 1000 bootstrap replicates [39], and visualized with iTOL [40].

### 2.4 Plant material and treatments

#### 2.4.1 Mutants and seed sterilization

Seeds of *Solanum lycopersicum* M82 were surface-sterilized with a sterilization solution (2.5% sodium hypochloride, 0.1% Tween-20) for 20 minutes. Seeds of *Arabidopsis thaliana* Col-0 were surface-sterilized with 70% ethanol and 0.05% Triton-X100 solution for 3 minutes. Immediately after, the seeds were washed with sterile distilled water five times. Tomato (*Slclv1-*a2, *Slbam1-*a1, *Slbam4-*a2 and *Slclv2-5*) and Arabidopsis (*Atcrn-10*) mutants are CRISPR-mediated mutants previously described [25, 41].

#### 2.4.2 Root assays

*S. lycopersicum* sterilized seeds were placed on 24 cm square plates containing 1µM of the indicated SlCLE peptide. After 2 days in the dark, plates were placed vertically in 16h light / 26 °C – 8h dark / 24 °C cycles for a week. *A. thaliana* sterilized seeds were grown onto 12cm square plates containing 50nM of indicated AtCLE peptides. After 2 days in the dark at 4 °C, plates were placed vertically in 16h light– 8h dark cycles at 22 °C for a week. The plates were scanned at high resolution, and primary root length was measured with the “simple neurite tracer” tool on Fiji (www.imagej.net). All CLE peptides are synthetic un-modified peptides at >75% purity (www.genscript.com) solubilized in water at 10mM stock concentration.

#### 2.4.3 Tomato drought stress assay

The assay was modified from a published protocol of hydroponically grown tomato [42]. In brief, sterilized tomato seeds were placed on moistened blotting paper and kept in dark at 26°C for 3 days. Germinated seeds were placed on Eppendorf-type tubes with cut end filled with 0.6% water-agar in 16h light / 26 °C – 8h dark / 24 °C cycles and high humidity environment for one week. Then, the seedlings were transferred to hydroponics containers, in which the roots grow in an oxygenated Hoagland solution in darkness. The nutritive solution was renewed every week. After 3 weeks, one day after replacing the nutrient solution, drought stress was induced with a fresh solution supplemented with 15% PEG-6000. Three different containers were used for the experiments generating each 2 biological replicates. Each biological replicate is a pool of 2 to 3 plants from the same container. The root samples contain all the root system coming out of the Eppendorf. The shoot samples contain all the leaves and around 5 cm of stem harboring the shoot apical meristem, thus these samples do not contain the main stem which has been strongly lignified.

### 2.5 Quantitative RT-PCR of tomato CLE genes

Plant tissues were rapidly shock frozen in liquid nitrogen. Frozen samples were grinded using mortar and pestle. Total RNA was extracted using the Spectrum Plant Total RNA kit (Sigma). The remaining DNA was eliminated by DNAse I treatment (Jena-Bioscience) and with a 2M LiCl precipitation. The absence of the genomic DNA in the RNA samples was tested by PCR. cDNA synthesis was performed using the SensiFAST cDNA synthesis kit (meridian). Quantitative PCRs were performed using Fast Start Universal SYBR-green Master (Roche), with primers indicated in Supplemental Table 1. The thermal cycler (Mic qPCR Cycler, biomolecular systems) conditions were: 95°C 2 min,45 cycles of 95°C 15s, 58 °C 10s, 60 °C 50s, followed by a dissociation curve analysis. The expression level was normalized to Actin on 6 biological replicates.

### 2.6 Microscopy

About 1 cm of the primary root tips of one-week-old tomato seedlings were fixed with 4% paraformaldehyde in a 1xPBS solution for a minimum of 6 hours. After 2 washes in 1xPBS, the samples were cleared in a ClearSee solution [43] for one week. Subsequently, to visualize the cell walls, the calcofluor white staining was performed with 0,02% calcofluor-white dissolved in the ClearSee solution for 2 days, followed by two washing steps with ClearSee. Samples were incubated in ClearSee solution for a minimum of 2 weeks before imaging. Images were taken with a confocal laser scanning microscope (Leica SP5). The calcofluor-white stained cell walls were excited at 405 nm and emitted light detected at 415-500nm. These images were used to quantify root width in the differentiation zone, columella length and cell number in Fiji (www.imagej.net).

### 2.7 Statistics

Statistical analysis was performed using Rstudio (www.rstudio.com) after log transformation of the data. Statistical significance was analyzed by ANOVA, and followed by a post-hoc Tukey test to determine the different statistical groups.

## 3. Results

### 3.1 Identification of 37 new *SlCLE* genes

Motivated by uncovering the role of CLE peptides in tomato roots, we performed a re-analysis of the CLE peptide-encoding genes. The first genome-wide analysis in the tomato genome revealed only fifteen *SlCLE* genes [23] and further attempts failed to uncover additional genes [21, 22]. In our study, we applied a combined bioinformatic approach to identify additional *SlCLE* genes using the most recent versions of the tomato reference genome SL3.0 and SL4.0 [44]. Firstly, we performed an iterative tBLASTn search on the full genome which revealed forty *CLE* genes (Figure 1B). Secondly, we applied a Hidden-Markov-Model, that resulted in forty-seven *CLE* genes. Remarkably, for a total of 52 *CLE* genes in tomato, fifteen were uncovered by only one of the two approaches. This indicates that the combination of these two methods can be instrumental to find short genes like *CLEs* in plant genomes.

**Figure 1.**
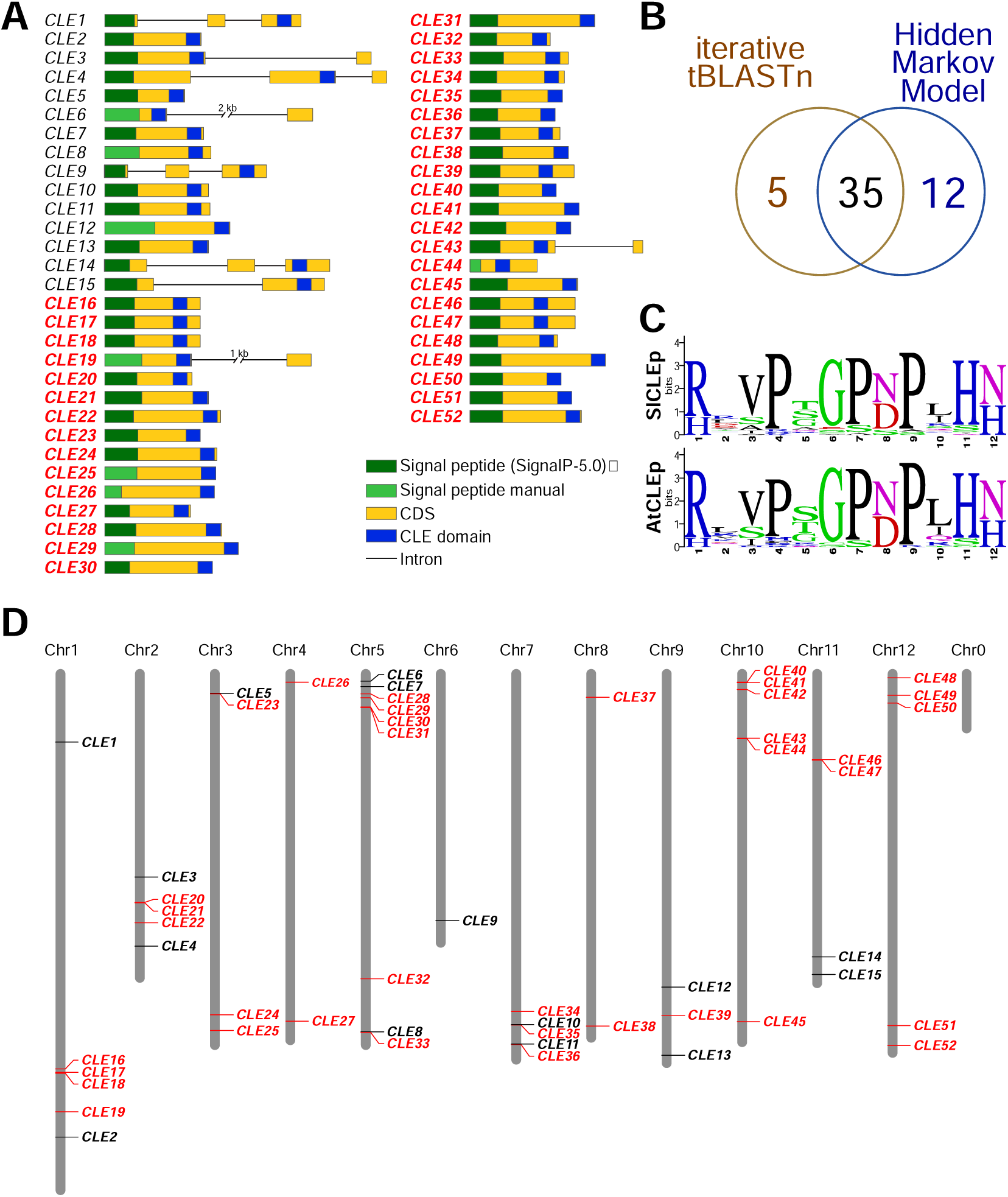
*CLE* genes identified in the tomato genome. **A**. The gene structure of the tomato *CLE* genes. Gene structures in dashed lines indicate the *SlCLE* genes for which no proof of expression was found in root, shoot and fruit tissues. **B**. Venn-diagramm showing the number of *CLE* genes identified by each method. **C**. Sequence logo of the conserved CLE domain in tomato and Arabidopsis using WebLogo (https://weblogo.berkeley.edu/logo.cgi) (reference). The height of the bars represents the conservation value of each amino acid at the given position. **D**. The chromosomal location of the tomato *CLE* genes.

**Figure 2.**
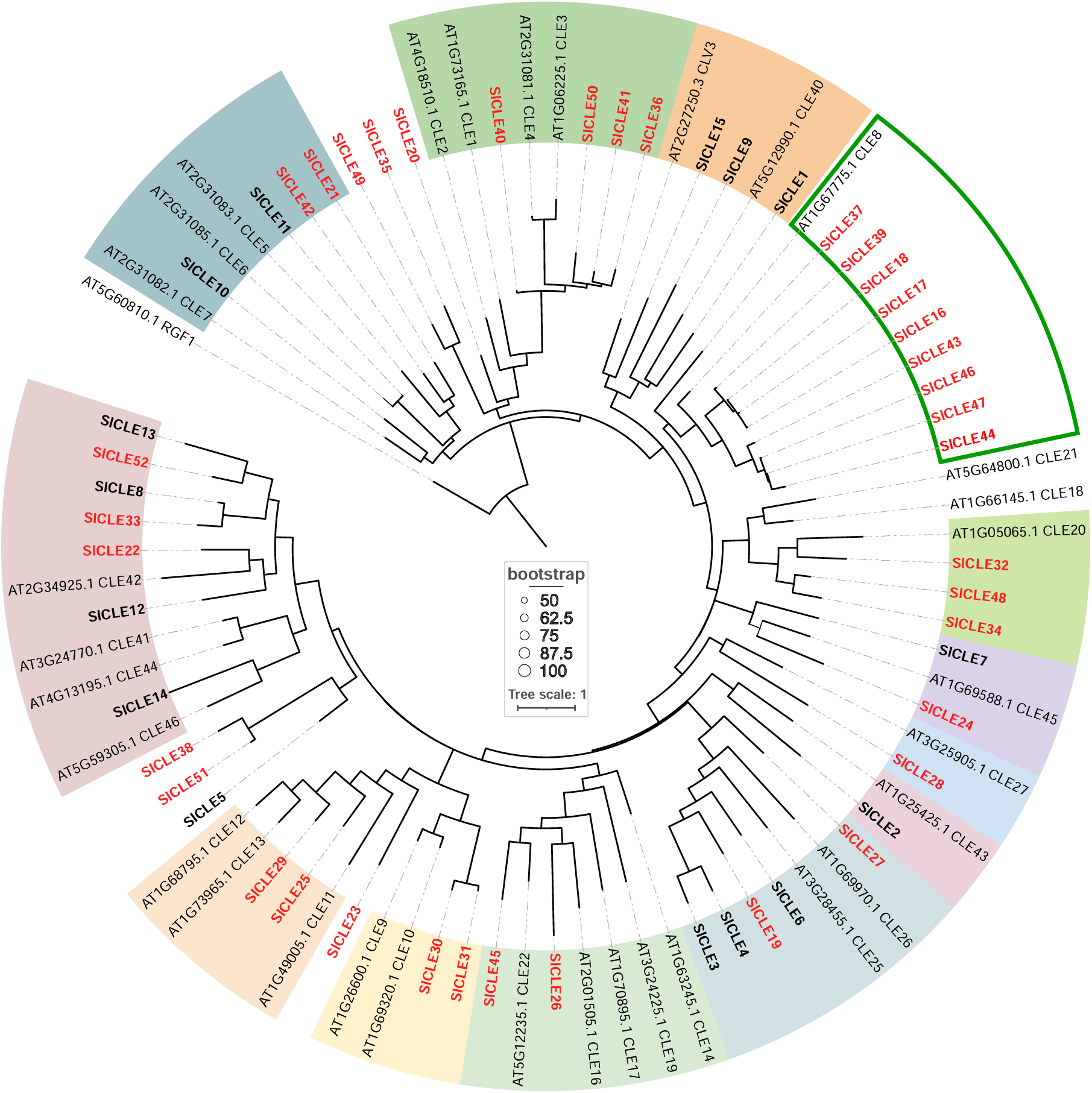
The phylogenetic tree of the full-length CLE proteins from tomato and Arabidopsis. The groups of proteins sharing high similarity (clusters) are highlighted by background colors. The names in red indicate the new *CLE* genes uncovered in this study. Nodes supported by bootstrap values superior to 50 are indicated by dots of size proportional to the bootstrap values.

We mapped the *SlCLE* genes on tomato’s chromosomes (Figure 1D). We numbered the identified *SlCLE* genes as follows: the previously reported fifteen genes are numbered *SlCLE1-SlCLE15*. The newly identified genes (*SlCLE16* to *SlCLE52*) are numbered according to their chromosomal location, starting from chromosome 1 (Figure 1D). *SlCLE* are diversely present on all 12 chromosomes in tomato, from a single gene on chromosome 6 to up to nine genes on chromosome 5. The fact that several *SlCLE* genes are located in high proximity with each other’s, forming gene clusters, and showing high sequence similarity, suggest that they arise from tandem gene duplication events. To investigate the gene structure diversity of tomato *CLE*s, the exon-intron composition was predicted based on sequence homologies (Figure 1A). In addition, we used publicly available RNAseq datasets [32-34, 45, 46], from root, shoot and fruit samples, to support these gene structure predictions. Reads were mapped on the anticipated coding region of 28 *CLE* genes out of the 37 newly uncovered loci (Figure 1A). Overall, the tomato *SlCLE*s have a single CLE domain in the 3’ of the coding region and do not include any intron (Figure 1A). In the case of *SlCLE31*, an insertion of a single nucleotide in the tomato genome SL4.0, which is not present in the version SL3.0, creates a frameshift in the CDS suggesting that it is a pseudogene. However, Sanger sequencing of this particular locus confirmed the correctness of the sequence in the SL3.0 genome. Furthermore, to evaluate to what extend the CLE motif is conserved between Arabidopsis and tomato, we created sequence logos (Figure 1C). The CLE domain is extremely well conserved, including the prolines at positions 4, 6, and 9, as well as the arginine at position 1, glycine at positions 6, and histidine-asparagine/histidine at positions 11-12.

In parallel, we searched for the CLE receptor genes, namely *CLV1, BAM1, BAM2, BAM3*, and *PXY*. As previously reported in tomato [25], we found one copy of *SlCLV1*, four *SlBAM* homologs, two *SlPXY*-like genes, one *SlPXL1*, and one *SlPXL2* (Supplemental Figure 2).

Tomato (*Solanum lycopersicum*) and potato (*Solanum tuberosum*) belong to the same family and the *CLE* genes in both share high gene sequence similarities. At the time our bioinformatics analysis was fully accomplished and the functional analysis of selected new *SlCLE* genes was still ongoing, a study reporting about 41 *CLE* genes in potato was published [20]. This study came up with a similar strategy for searching short genes in the potato genome. We used the advantage of this list of potato CLE genes to perform a phylogenetic analysis with tomato genes identified in present study (Supplemental Figure 1). Except for *StCLE2* and *StCLE5*, we found orthologous for all the other CLE genes in the potato genome, which indicates that both studies identified most of the CLE genes.

### 3.2. Expression analysis of *SlCLE*s and their diversification

Next, we wanted to confirm, that the identified genes are expressed in tomato based on the publicly available RNAseq datasets [32-34, 45, 46]. Remarkably, the majority of *SlCLEs* shows predominant expression in root tissues, while some are shoot-specific or evenly expressed in both (Figure 3A). We could observe, that *SlCLE20, SlCLE2, SlCLE40* and *SlCLE41*, for example, are root-specific genes and their expression increases with the age of the plant. Using qPCR, we could detect that *SlCLE5, SlCLE21, SlCLE40* have higher expression in the root tissues, while *SlCLE13, SlCLE32, SlCLE45*, and *SlCLE52* are more expressed in the shoot tissues (Figure 3B).

**Figure 3.**
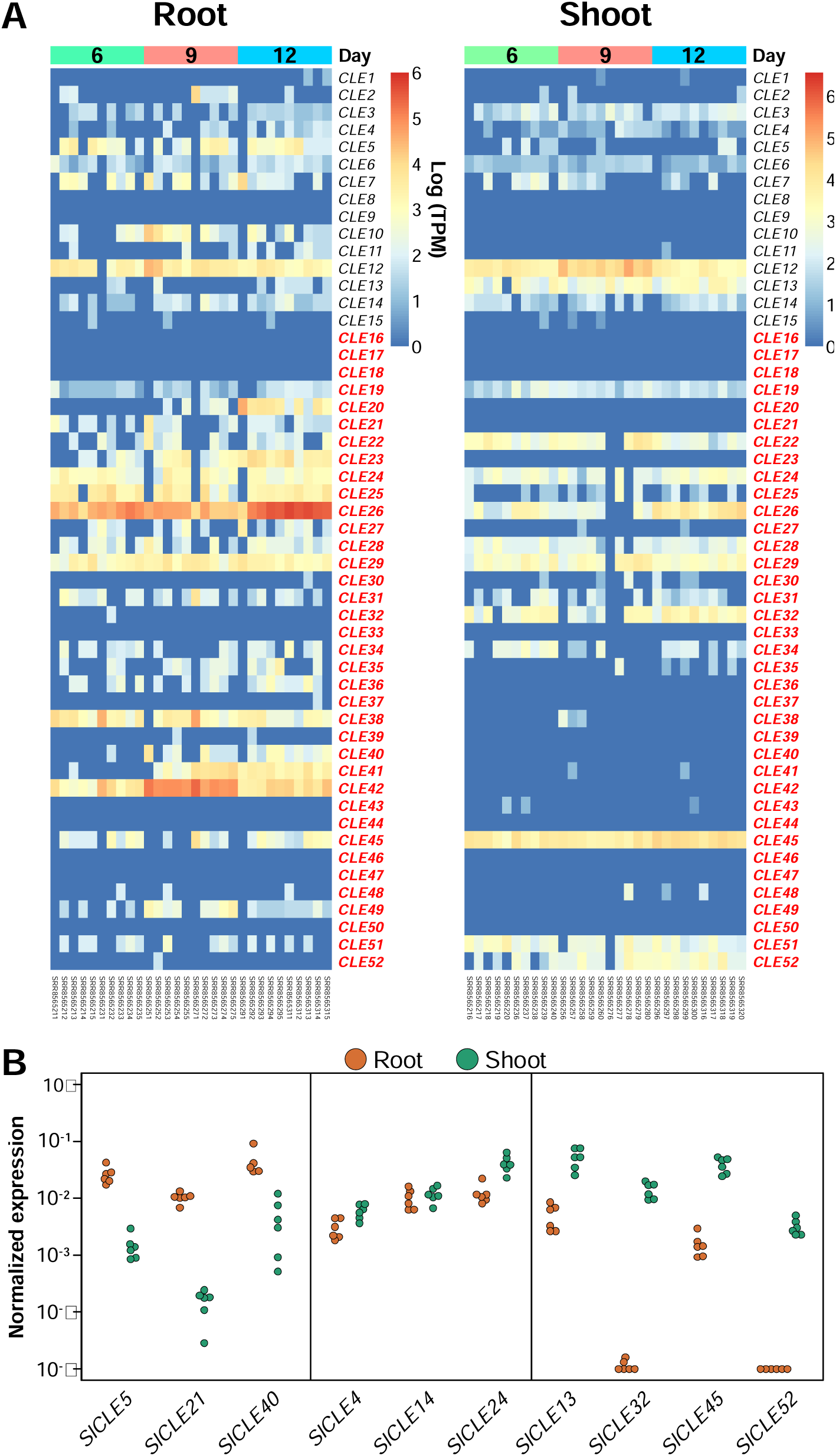
Expression analysis of *SlCLE* genes in root and shoot tissues [34]. **A**.Heatmaps of log(TPM) of tomato *CLE* genes in the root (left) and the shoot (right) at 6, 9, and 12 days after plantation from tomato grown in pots. **B**. Expression of selected *SlCLE*s in root and shoot tissues by qPCR from 3 weeks old tomato plant grown in hydroponic conditions.

To explore the diversification of the tomato *CLE* genes, we created a phylogenetic tree of the full-length proteins from tomato and Arabidopsis (Figure 2). This analysis revealed gene sub-groups that are conserved in both plant species as well as showed unique genes, which could pinpoint *CLE* diversifications in tomato or losses in Arabidopsis, and define orthology. Interestingly, in tomato, we found nine homologs of Arabidopsis CLE8 (Figure 2). In Arabidopsis, this peptide is expressed and acts specifically during embryo and endosperm development [47], but the roles of the nine orthologs in tomato are yet to be uncovered. For phloem CLE peptides in Arabidopsis, which includes CLE25, CLE26, CLE45 we found seven orthologs in tomato, which also suggests the diversification of the phloem genes.

### 3.3 The expression of *SlCLEs* in the developing fruit and in water deficit stress

To test whether *SlCLE* genes are expressed during fruit development, we analyzed the dataset from [33]. In this study, wild type M82 and yellow-fruited *yft1* mutant fruits were sampled at different developmental time points, from 35 to 60 days-post-antherisation. Our analysis shows, that *SlCLE12, SlCLE30, SlCLE31, SlCLE34*, and *SlCLE38* are the most expressed in tomato fruits independently of the genotype, whereas *SlCLE5, SlCLE11, SLCLE51* expression is impaired in the yellow-fruited *yft1* mutant (Supplemental Figure 3A). These results suggest, that *SlCLE* genes could play a role during tomato fruit ripening.

Numerous studies showed that CLE peptides mediate abiotic stress signals, for example, CLE25 peptide in Arabidopsis was shown to be induced during dehydration, moving from root to shoot as a mobile signal, triggering ABA biosynthesis and stomatal closure [5]. Therefore, we wanted to test, whether some *SlCLE* genes are up-regulated under drought stress conditions. First, we analyzed the dataset published previously and could find several genes that are specifically expressed under drought in the tomato leaves (Supplemental Figure 3B) [32]. *SlCLE1, SlCLE12, SlCLE32, SlCLE45* and *SlCLE52* showed an increased expression (Supplemental Figure 3B), suggesting that they could be involved in adaptive responses to water deficit We wanted to test, whether these genes are quickly up-regulated, also following short osmotic stress. To this end, hydroponically grown tomato plants were treated with a 15% PEG6000 solution for one hour, and roots and shoots samples were collected separately. Since Dehydrins (DHN) play a key role in plant response and adaptation to water deficit conditions and are accumulated during drought stress, we used the *SlDehydrin (SlDHN)* (*Solyc02g084850*) expression as a control to monitor the effect of water deficit in our experiment. After one hour, *SlDHN* was strongly upregulated both in root and shoot tissues of treated tomato plants (Supplemental Figure 3C). However, we could not detect a significant induction for those *SlCLE* genes (Supplemental Figure 3C). Next, we wanted to test, whether similarly to Arabidopsis, the tomato orthologs of *AtCLE25* are upregulated in roots to mediate a dehydration response like it has been demonstrated in Takahashi et al 2018 [5]. We could not detect any significant induction in *AtCLE25* orthologs in tomato under this short osmotic stress (Supplemental Figure 3D). One possibility is that our experimental settings were not enough to trigger similar osmotic stress like reported in [5] and [32]. Another possibility is that in tomato, none of the *AtCLE25* orthologs are involved in mediating drought responses.

### 3.4 A conserved effect of *SlCLE* peptides on root apical meristem

To test whether the predicted CLE peptides are biologically active, we performed root growth assays with commercially synthesized unmodified peptides. It has been shown, that in Arabidopsis 20 out of 32 peptides affect the primary root growth, leading to root meristem arrest [9, 48]. To study the activity of orthologous CLE peptides, we tested their capacity to inhibit root growth. For this purpose, we selected CLE peptides from different subgroups and well-supported orthologous genes in tomato and Arabidopsis, which were known to have different potency in arresting the root meristem in Arabidopsis. For example, the treatment with AtCLV3, AtCLE25, and AtCLE45 peptides at 50nM triggers a strong reduction of the primary root length (Figure 4A right side), and this response depends on the pseudo-kinase CORYNE and the receptor-like protein CLAVATA2 [9]. However, the AtCLE9/10 and AtCLE22 peptides had a much smaller root growth inhibition effect in the wild-type.

**Figure 4.**
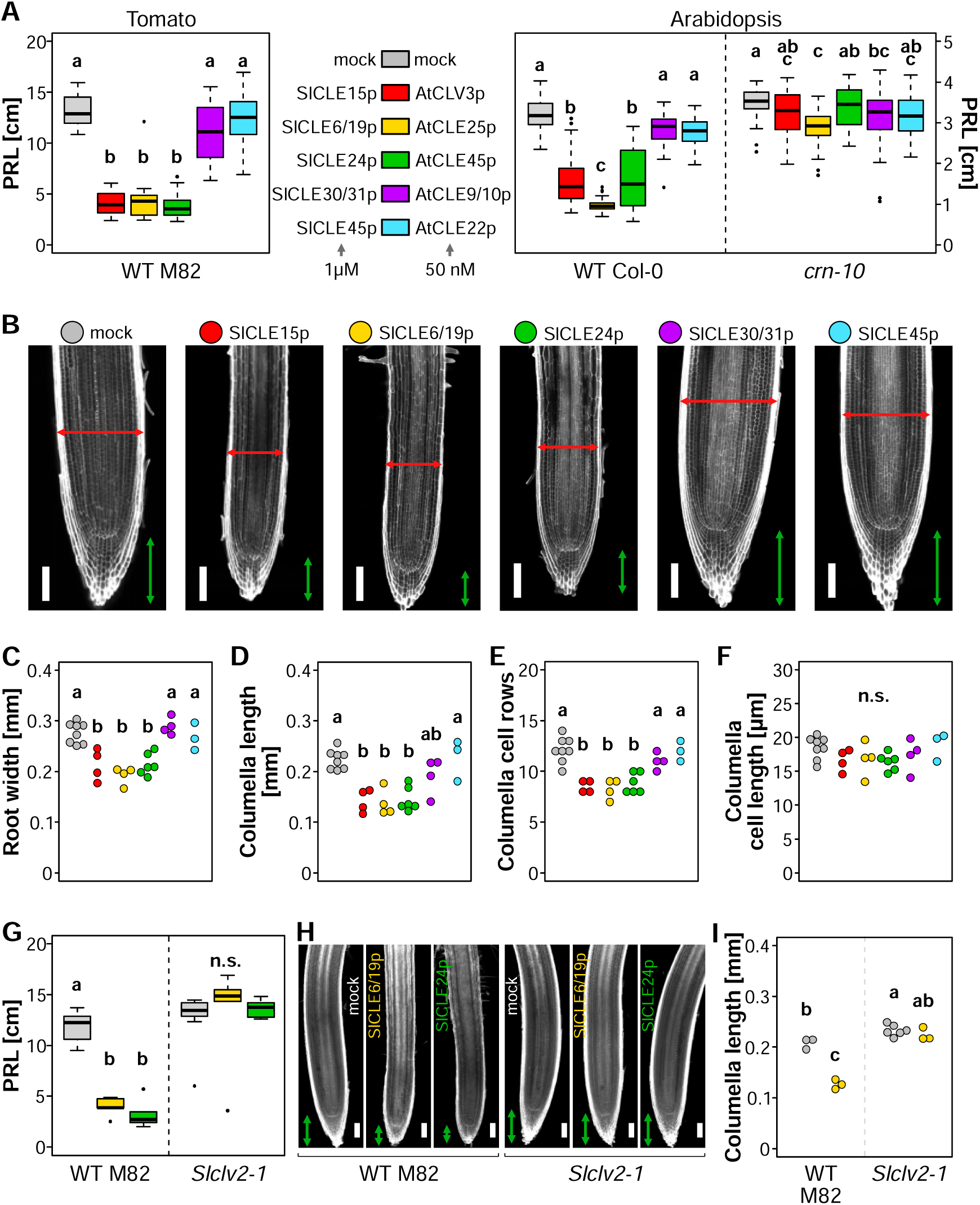
Functional conservation of root active CLE peptides in tomato and Arabidopsis. **A**.Effect of Arabidopsis and tomato orthologous CLE peptides on the primary root length (PRL). **B**. Representative confocal images of tomato primary root tips grown on mock or indicated SlCLE peptide containing medium. The cell walls are stained with calcofluor white. Red and green arrows indicate root width columella length, respectively **C-F**. Quantification of the indicated root tip morphology characteristics from images show in B. Letters indicate different statistical groups (ANOVA, followed by post-hoc Tukey test). **G**. Primary root length of wild type and *Slclv2* mutant grown in presence of SlCLE6/19p and SlCLE24p. **H**. Representative confocal images of wild type and *Slclv2* primary root tips grown on mock or indicated SlCLE peptide containing medium. The scale bars in **B** and **H** correspond to 100µm.

Because tomato roots are much thicker, with a diameter of about 320 µm at the tip instead of approximately 120 µm in Arabidopsis, tomato root tip is less sensitive to external application of CLE peptides. Therefore, we applied SlCLE peptides at a concentration of 1 micromolar. We could observe, that in tomato roots, SlCLE15, SlCLE6/19 and SlCLE24 peptides (orthologous of AtCLV3, AtCLE25, and AtCLE45, respectively) led to a strong reduction of the primary root growth (Figure 4A left side). In contrast, SlCLE30/31 and SlCLE45 peptides (orthologous of AtCLE9/10 and AtCLE22, respectively) treatment did not trigger a significant reduction of the primary root length. This result indicates that the amino-acid composition of CLE peptides is important for their biological activity in the root; but, more importantly, that there is a conservation of the biological activity of these CLE peptides between Arabidopsis and tomato, two evolutionary separated species.

To have more insight into the effect of SlCLEs on tomato roots, we analyzed the morphology of the root tips (Figure 4B). We observed in three treatments (SlCLE15, SlCLE19, SlCLE24) not only the decreased root length but also the reduction in the root diameter and the columella length. We also looked at the number of columella layers and the columella cell length to understand whether the treatment affects cell division or cell elongation. Root-active SlCLE peptides treatment led to a reduction of columella layers, but not their average cell length, suggesting that cell division is primarily affected (Figure 4C-F). We can conclude, that the identified CLE peptides have a root inhibitory effect by restricting the meristematic cell divisions.

Next, we asked whether this inhibitory effect on the root is mediated by orthologous receptor-like kinases? To answer this question, we tested the loss-of-function mutants *clv1 bam1 bam4* and *clv2* [25] for their root sensitivity to SlCLE peptides. The mutant *clv1 bam1 bam4* showed a strong sensitivity to SlCLE24 peptide (Supplemental Figure 4). However, *clv2* roots were absolutely blind to the high concentrations of the peptide in the media, strongly suggesting that this response is *SlCLV2*-dependent. This result reinforces the claim, that CLE peptides have a conserved root activity across plant species and that the perception mechanism is similar.

## 4. Discussion

Signaling mediated by CLE peptides evolved gradually in all land plant lineages [49]. The precise control of the shoot apical meristem stem cell niche by CLV3-CLV1 module is the most ancient pathway, whereas additional CLE genes and additional receptor complex components evolved later, with establishing vascular plants [49]. It seems, that the possible ancestral function of CLV3-like peptides was to suppress the proliferation of the shoot apical meristem in early land plants (bryophytes). Pseudo-kinase CRN and receptor-like protein CLV2 create a complex and localize to the plasma membrane to mediate CLE signals. CRN, CLV2 and additional CLE genes appeared later in the vascular plants. To uncover the full repertoire and to understand the function of these “later” CLE genes in tomato, we performed a re-analysis. We have discovered thirty-seven new *SlCLE* genes and we could show that they are expressed and likely result in active peptides.

Strikingly, the perception of SlCLEs in the tomato root meristem is highly conserved and requires SlCLV2 receptor-like protein. The loss-of-function *Slclv2* mutant is fully insensitive to the peptide treatment. In tomato wild type roots, the root-active synthetic peptides (orthologs of Arabidopsis CLV3, CLE25, and CLE45) inhibited the cell division rate in the columella and vascular tissues. It has to be carefully analyzed whether such orthologs share the tissue-specific pattern and the endogenous biological function.

It has been shown, that SlCLV3 and SlCLE9 undergo arabynosylation and are active at 60 nM concentration while glycosylated and are not active at this concentration if the peptide is unmodified [24]. Moreover, it has been demonstrated that the biological activity of Arabidopsis CLV3 gradually increases in mono-, di- and triarabinosylated CLV3 glycopeptides, becoming equally active with non-modified peptide at 1 µM concentration [50]. The synthesizing of the complex arabinose chain is technically difficult and only a few laboratories in the world established such synthesis [50]. It is plausible, that the effect of glycosylated SlCLEs on the root meristem will be visible at a much lower concentration.

Our study showed that the root sensitivity to SlCLE treatment was not dependent on *SlBAM1, SlBAM4* or *SlCLV1*. This result suggests that in tomato roots additional *SlBAM*s can act redundantly in perceiving the SlCLEs. It will be interesting to test the combinations of *SlBAM1* and *SlBAM3* loss-of-function, and *SlBAM1* and *SlBAM2* loss-of-function, for example. The receptor-like kinase SlBAM4 is present in tomato, but not in Arabidopsis genome and the *Slbam4* loss-of-function mutant remains sensitive to the peptide treatment. This receptor-like kinase-encoding gene was lost in the Arabidopsis genome during the evolution and the function of this gene in tomato has not yet been studied.

The phylogenetic analysis and the clustering of the *SlCLE* genes with the Arabidopsis orthologs allowed us to detect very interesting diversification events. For example, we found nine orthologs of *AtCLE8* that was shown to control the embryo development [47]. The role of these genes remains to be uncovered. In conclusion, our work draws a more precise picture of the components of CLE signaling in this fleshy fruit crop plant paving a path for new discoveries.

## Supporting information

The list of the SlCLEs identified in this study

Primers used in this study

## Author contributions

The manuscript was conceptualized by OH and SC. The bioinformatic analysis was performed by SC and LF; the biological experiments and the preparation of the figures were performed by SC. The first version of the manuscript was written by OH. All authors contributed to the editing of the final version of the manuscript.

## Acknowledgments

We thank Prof. Yalovsky (Tel Aviv University, Israel) for sharing the tomato wild type (M-82) seeds initially. We thank Prof. Lippman (Cold Spring Harbor Laboratory, US) for sharing with us the tomato receptors mutants previously published in [25].

The research on tomato and Arabidopsis CLE signaling is funded by Ambizione SNSF grant (PZ00P3_179745) to OH, COST SNSF Grant (IZCOZ0_189892) to OH, and additional funding provided by the Department of Biology to OH.

## Supplemental Figures and Tables

**Supplemental Figure 1. Sequence homology between tomato and potato CLE proteins**. Phylogenetic tree of full-length CLE proteins from tomato (red), potato (blue), *Arabidopsis thaliana, Medicago truncatula, Brachypodium distachyon*. Nodes supported by bootstrap values superior to 50 are indicated by dots of size proportional to the bootstrap values.

**Supplemental Figure 2. Phylogenetic analysis of CLE receptors**.The receptor genes from *Arabidopsis thaliana* and tomato are in bold black and red, respectively.

**Supplemental Figure 3. Expression of tomato *CLE* genes in the developing fruit and following drought stress. A**. Heatmaps of log (TPM) of *SlCLE* genes in the developing fruit at 35, 47, 54, and 60 days post antherisation (DPA), in wilt-type and *yellow fruit tomato1* mutant. Genes in red indicate *SlCLE* which are upregulated during fruit maturation in a *yft1* dependent fashion. **B**.Heatmaps of log (TPM) of *SlCLE* genes in plant leaves (M82 or a drought tolerant introgression line IL9-1) after a 10 days drought stress in pots. Genes in red indicate *SlCLE* which are upregulated by the drought treatment. **C**. Expression analysis by qPCR of selected *SlCLE* genes on root and shoot samples from 3 weeks old plants grown in hydroponic after 1 hour 15% PEG6000 treatment. **D**. Expression analysis of the orthologs of *AtCLE25* in response to osmotic stress induced by the PEG treatment.

**Supplemental Figure 4**.Effect of SlCLE24 peptide on the primary root length (PRL) in wild type (M82) and *clv1bam1bam4* mutant.

**Supplemental Table 1. Primers used in this study**.

**Supplemental Table 2. Sequences and chromosome locations of the *SlCLE*s**

