## Supplemental figures for "New insights into tomato CLE peptide repertoire and perception mechanisms"

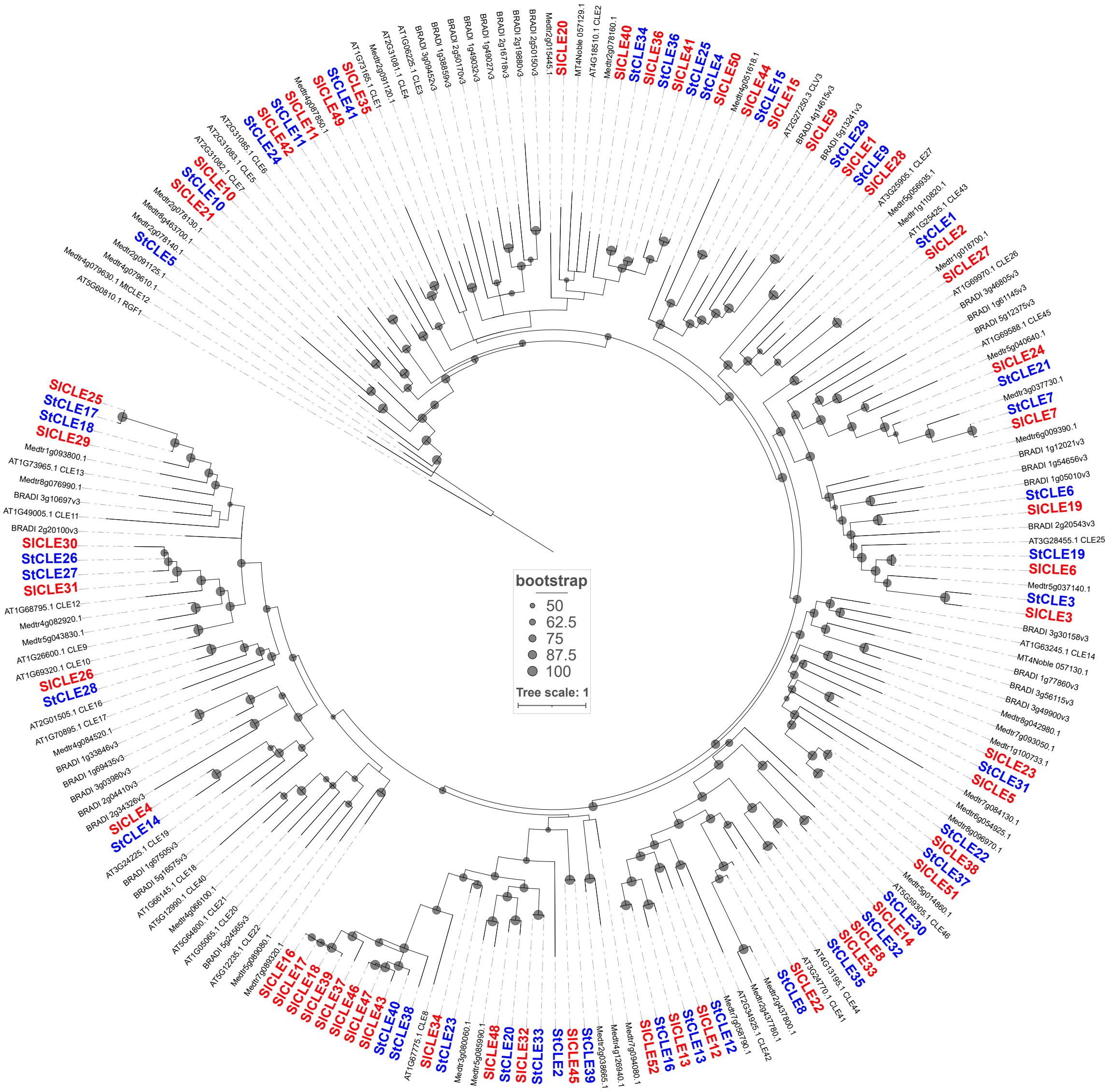


**Supplemental Figure 1. Sequence homology between tomato and potato CLE proteins.** Phylogenetic tree of full-length CLE proteins from tomato (red), potato (blue), *Arabidopsis thaliana*, *Medicago truncatula*, *Brachypodium distachyon*. Nodes supported by bootstrap values superior to 50 are indicated by dots of size proportional to the bootstrap values.

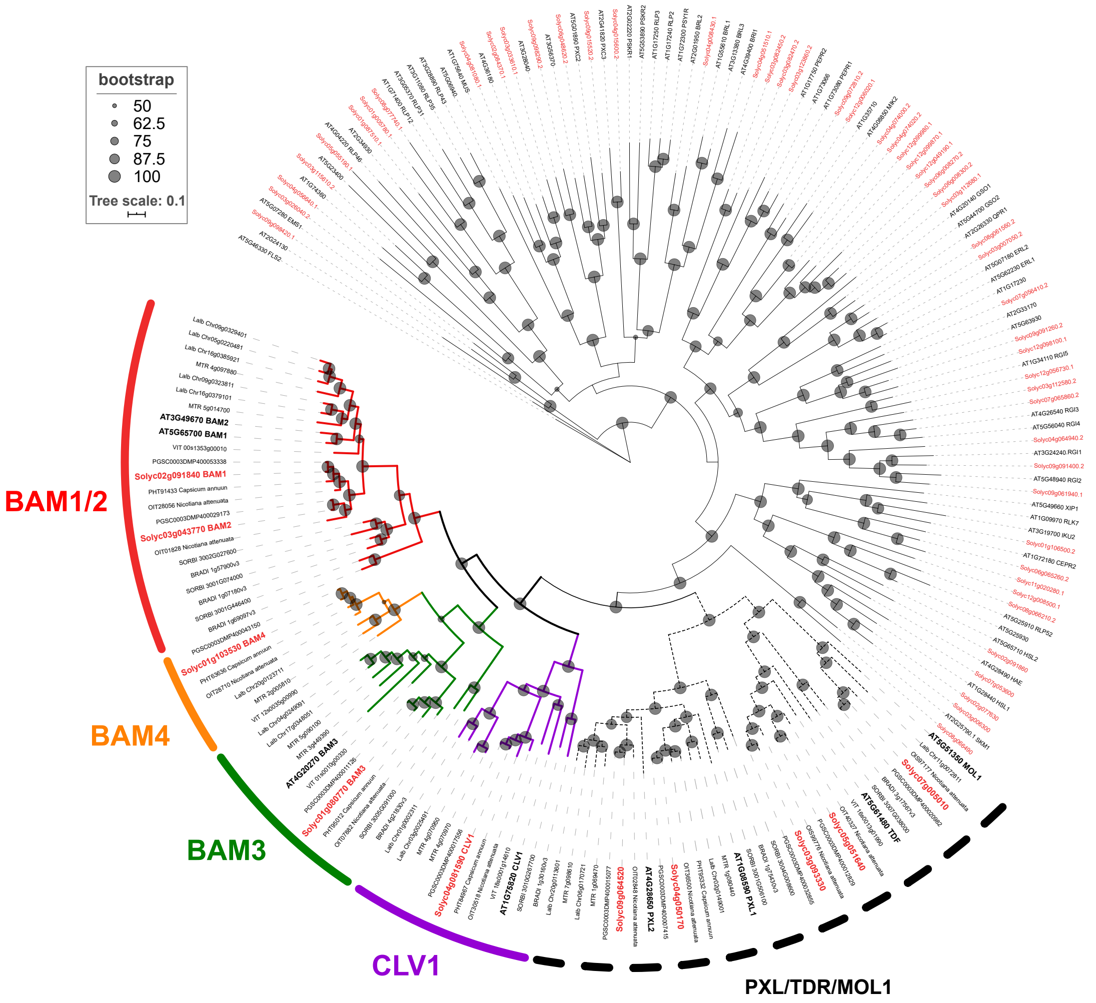


**Supplemental Figure 2. Phylogenetic analysis of CLE receptors**. The receptor genes from *Arabidopsis thaliana* and tomato are in bold black and red, respectively.

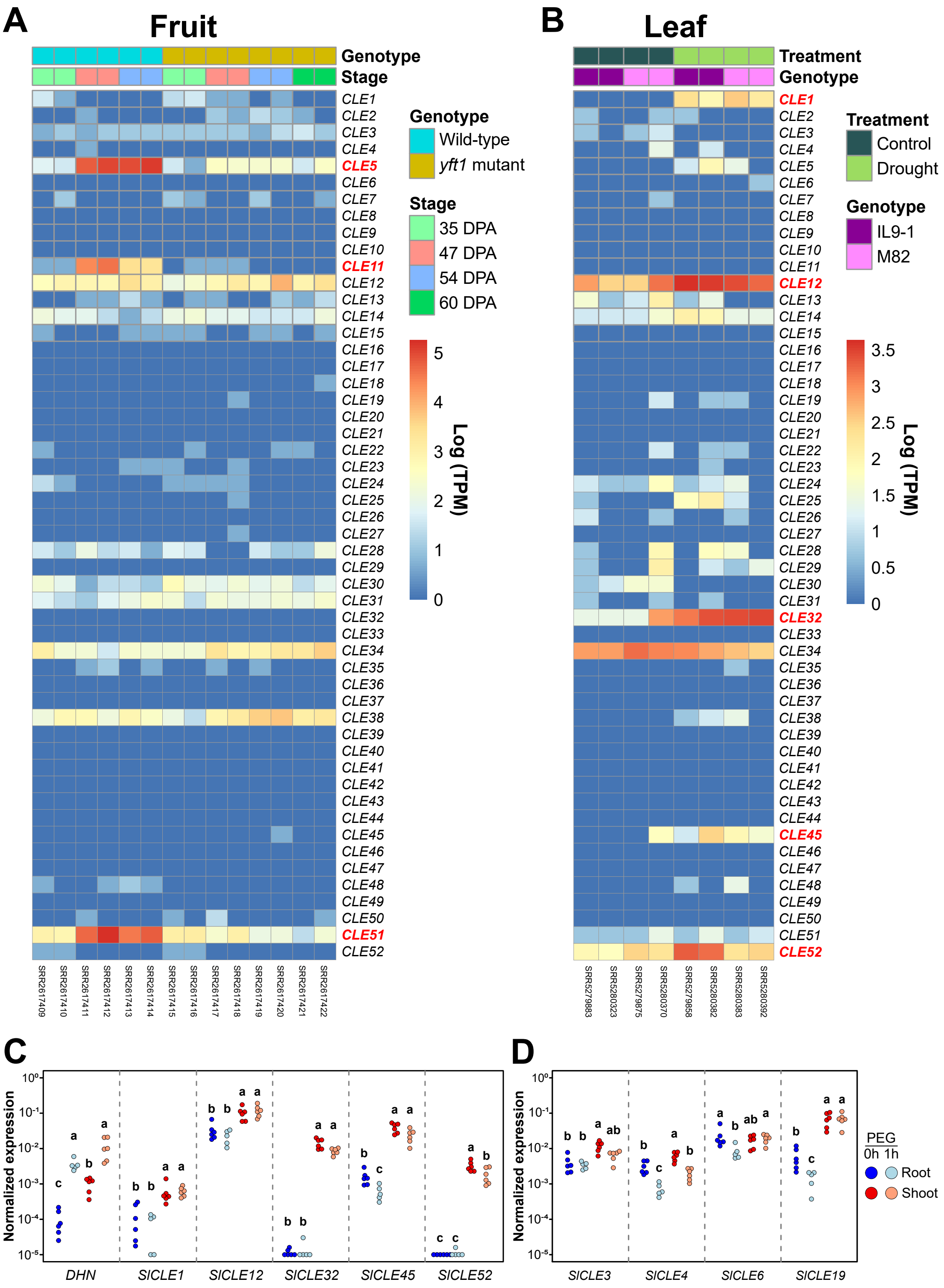


**Supplemental Figure 3. Expression of tomato *CLE* genes in the developing fruit and following drought stress. A.** Heatmaps of log (TPM) of *SlCLE* genes in the developing fruit at 35, 47, 54, and 60 days post antherisation (DPA), in wilt-type and *yellow fruit tomato1*

mutant. Genes in red indicate *SlCLE* which are upregulated during fruit maturation in a *yft1* dependent fashion. **B.**Heatmaps of log (TPM) of *SlCLE* genes in plant leaves (M82 or a drought tolerant introgression line IL9-1) after a 10 days drought stress in pots. Genes in red indicate *SlCLE* which are upregulated by the drought treatment. **C.** Expression analysis by qPCR of selected *SlCLE* genes on root and shoot samples from 3 weeks old plants grown in hydroponic after 1 hour 15% PEG6000 treatment. **D.** Expression analysis of the orthologs of *AtCLE25* in response to osmotic stress induced by the PEG treatment.

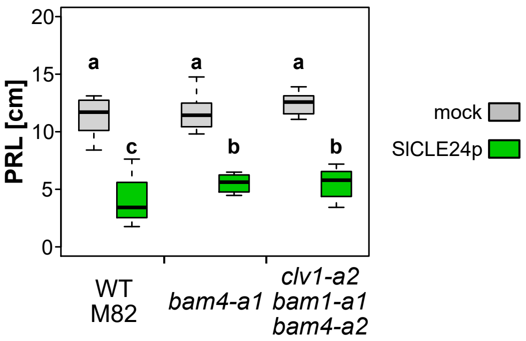


**Supplemental Figure 4.** Effect of SlCLE24 peptide on the primary root length (PRL) in wild type (M82) and *clv1bam1bam4* mutant.
