## Supplementary material for "New insights into tomato CLE peptide repertoire and perception mechanisms": Primers used in this study

Table1. qPCR primers.

| Number | Gene | Sequence | Reference |
|---|---|---|---|
| F01 | *SlActin* | GGTCCTCTTCCAGCCATCC | Zhang et al., BMC 2014 |
| F02 | | CCACTGAGCACAATGTTACCG | |
| H11 | *SlDHN* | CACCATGAGGGGCAACAGCA | Kissoudis et al., Front. Plant Sci. 2017 |
| H12 | | TCACCTTCATGTTGTCCAGGCATC | |
| E52 | *SlCLE1* | TGGTGTCTTTAAGAACTTTTGCTG | Zhang et al., BMC 2014 |
| E53 | | CTCTTTATCTGGAAAATCCCCTT | |
| E56 | *SlCLE3* | CTGCTGAGATTTTAGTAAAGCCTG | Zhang et al., BMC 2014 |
| E57 | | GAATGCCTTTCTGTTTCTATTATCC | |
| F58 | *SlCLE4* | GGGAAGGGAACTAGGGCTGCCA | This study |
| F59 | | TGGCATTTGGTCCAGTAGGCACT | |
| E60 | *SlCLE5* | AACCTCCCACTTCATTACTTCTTC | Zhang et al., BMC 2014 |
| E61 | | ATGATCTGCAGCACCAGCAT | |
| E62 | *SlCLE6* | TGGAGGTGTTACAACAAAAATGA | Zhang et al., BMC 2014 |
| E63 | | GAACATGATGAGCACCACTTGA | |
| E74 | *SlCLE12* | TGATGGATATTGATCTCTTGTGGA | Zhang et al., BMC 2014 |
| E75 | | ATGAATGGTTGGGAACTGGAT | |
| E76 | *SlCLE13* | CAATATGCAAGTCCATCACAAAC | Zhang et al., BMC 2014 |
| E77 | | GCCTCCCCATAATATTTTCGA | |
| J08 | *SlCLE19* | CCTAATGGCCCAGACCCTAT | This study |
| J09 | | GGCTTGCCAAATTCTCCTTT | |
| F62 | *SlCLE21* | TCGTGGAGTCGAGAACATGAGGA | This study |
| F63 | | GGATCTGGTCCTCCAGGTGCAA | |
| F66 | *SlCLE24* | AAGGCTGCTGTCGTGCAAGACC | This study |
| F67 | | CCCTGCACACTTGATTTGGACTCGC | |
| F78 | *SlCLE32* | ACCTCTCCAAGAGTTCATGTCATCCA | This study |
| F79 | | AATATGACACTCTCTTTTCGTGGCGA | |
| G09 | *SlCLE40* | TGTCCTCCCTCCGAAAGTCGTCC | This study |
| G10 | | TATCGTTTCCCCTCTCCTCCGCG | |
| G15 | *SlCLE45* | TGAAAACCCCATGAGCCATGACT | This study |
| G16 | | TCCTCTTGAGAAGAAGCTTCCTTTGT | |
| G25 | *SlCLE52* | ACCACGACCACCACTACTGTCA | This study |
| G26 | | GCAGCCACCCCATATTGCCCTC | |
